# PATAN-domain response regulators interact with the Type IV pilus motor to control phototactic orientation in the cyanobacterium *Synechocystis* sp. PCC 6803

**DOI:** 10.1101/2021.06.11.447907

**Authors:** Yu Han, Annik Jakob, Sophia Engel, Annegret Wilde, Nils Schuergers

**Affiliations:** Molecular Genetics, Institute of Biology III, University of Freiburg, 79104 Freiburg, Germany

**Author notes:** corresponding author <u>**Email:**</u>.

**Keywords:** Phototaxis, chemotaxis, type IV pili, PATAN, CheY

## Abstract

Many prokaryotes show complex behaviors that require the intricate spatial and temporal organization of cellular protein machineries, leading to asymmetrical protein distribution and cell polarity. One such behavior is cyanobacterial phototaxis which relies on the dynamic localization of the Type IV pilus motor proteins in response to light. In the cyanobacterium *Synechocystis*, various signaling systems encompassing chemotaxis-related CheY- and PatA-like response regulators are critical players in switching between positive and negative phototaxis depending on the light intensity and wavelength. In this study, we show that PatA-type regulators evolved from chemosensory systems. Using fluorescence microscopy and yeast-two-hybrid analysis, we demonstrate that they localize to the inner membrane, where they interact with the N-terminal cytoplasmic domain of PilC and the pilus assembly ATPase PilB1. By separately expressing the subdomains of the response regulator PixE, we confirm that only the N-terminal PATAN domain interacts with PilB1, localizes to the membrane, and is sufficient to reverse phototactic orientation. These experiments established that the PATAN domain is the principal output domain of PatA-type regulators which we presume to modulate pilus extension by binding to the pilus motor components.

## Introduction

Phototaxis allows cyanobacteria to seek out optimal conditions for their photosynthetic lifestyle. Depending on the quality and intensity of illumination, single cells and whole colonies alike can move either towards (positive phototaxis) or away (negative phototaxis) from a light source (1, 2). This behavior requires the intricate spatial and temporal coordination of the motility machinery. The single-celled cyanobacterium *Synechocystis* sp. PCC 6803 (hereafter *Synechocystis*) is a model for studying the mechanisms of phototaxis control in cyanobacteria. During phototaxis, *Synechocystis* establishes directionality by focusing the incoming illumination within its lens-like cell body. The local excitation of receptors in the focal spot is thought to guide the asymmetrical distribution of PilB1, resulting in polar pilus activity and phototactic movement in regard to the light vector (3–5). The integration of different light signals governs phototactic orientation. Wild-type cells exhibit positive phototaxis towards wavelengths ranging from green to far-red, whereas blue or UV-light inhibit motility or induce negative phototaxis (1, 2, 6).

Like many bacteria, *Synechocystis* moves along solid surfaces using type IVa pili (TFP), protein filaments that also play a role in adhesion, aggregation, and DNA uptake (7–10). Studies on other bacteria demonstrated that TFP-mediated “twitching” motility relies on the sequential extension, surface adhesion, and retraction of the pilus filament to pull the cell forward (11). Translocation of the pilus through a complex that spans the cell envelope is driven by the inner membrane protein PilC that promotes the (dis-)assembly of the pilus that is composed of major and minor pilin subunits (12). Two antagonistically working motor ATPases can interact with PilC at the cytoplasmic face of the inner membrane to power cell movement (13, 14). In an ATP-driven process, PilB facilitates pilus assembly while PilT is required for pilus retraction (15). In *Synechocystis*, which encodes two PilB and PilT homologs, PilB1 and PilT1 fulfill these functions (7, 9, 16). The additional homologs of the motor ATPases, PilT2 and PilB2, have so far unknown functions.

Different signal transduction pathways regulate pilus assembly or control the reversal of cell polarity (i.e., the switch from positive to negative phototaxis). The most prevalent systems in cyanobacteria are homologous to the chemosensory systems that regulate flagellar rotation in many chemotactic bacteria. The sensor of such a canonical system is a methyl-accepting chemotaxis protein (MCP) that transduces a chemical cue to the histidine kinase CheA, which is linked to the MCP via CheW. Upon autophosphorylation of CheA, the phosphoryl group is transferred to the response regulator CheY that diffuses to the base of the flagellum and promotes a switch in the direction of flagellar rotation (17). Homologous systems in cyanobacteria typically harbor two CheY-like response regulators but miss the CheR and CheB proteins that modulate MCP methylation and temporal adaptation in canonical systems (18). *Synechocystis* encodes three of these systems, designated Tax1-3, for their role in phototaxis (19) (Fig. 1A). The Tax1 system with the blue/green absorbing cyanobacteriochrome photoreceptor PixJ1 is essential for positive phototaxis, and disruption of the response regulator genes *pixG* or *pixH* leads to negative phototaxis in response to white light (2, 19–21). The MCPs of the Tax2 and Tax3 systems lack a discernible photoreceptor domain, and it is unclear which signaling cues they perceive. While disruption of the Tax2 system does not affect motility (19), mutations in the Tax3 pathway lead to defects in pilus assembly that abrogate cell movement completely (19, 22, 23). Notably, the Tax3 system is linked to pilus assembly and/or retraction; insertional mutations in sequences encoding the second response regulator (*pilH*) or the phosphotransfer domain of the CheA homolog (*slr0073*) encoded by the *tax3* gene cluster cause a hyper-piliated phenotype, whereas insertions near the dimerization domain of CheA (*slr0332*), or within the MCP-homolog (*slr1044*) lead to a loss of TFP on the cell surface (22). Thus far, it is unresolved how the Tax1 and Tax3 signaling pathways transduce their respective signals to the motility machinery.

**Fig. 1:**
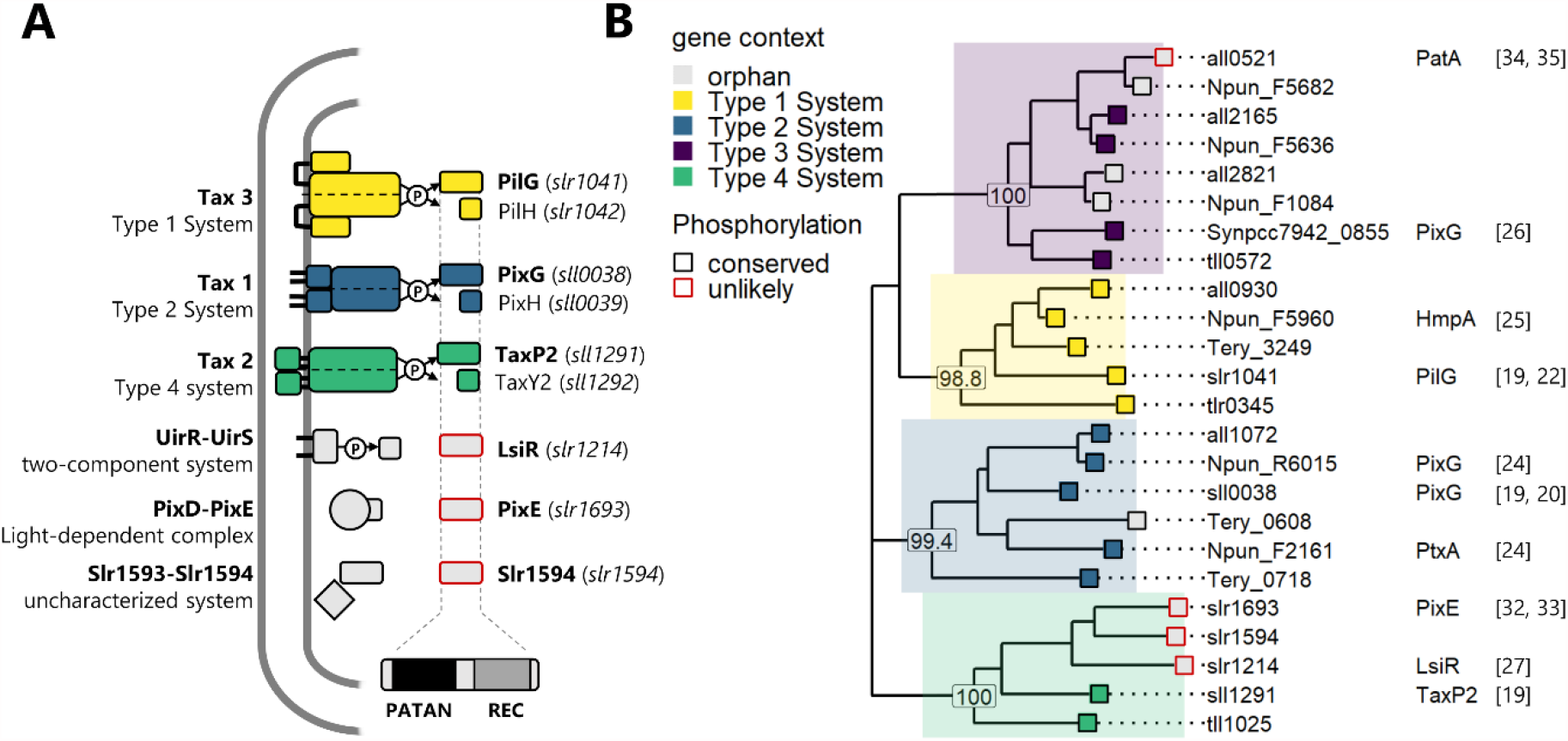
Cyanobacterial PatA-type response regulators originate from ancestral chemosensory systems. (A) *Synechocystis* harbors six PatA-type response regulators comprising an N-terminal PATAN domain and a C-terminal REC domain. Three of these are encoded upstream of a CheY homolog as part of the chemotaxis-like signaling systems Tax1-Tax3. The other three occur in the context of entirely unrelated signaling systems. (B) Maximum likelihood phylogenetic tree of PatA-type regulators identified from a sample set of cyanobacterial sequences aligned with MAFFT via Guidance2 with bootstrap probabilities calculated using 500 replicates shown for selected nodes. For PatA-homologs that are part of a chemosensory cluster, the type of the system as determined from the phylogeny of the associated CheA histidine kinases (see Fig. S1) is color-coded. The line color of the tip indicates whether conserved residues required for phosphorylation of the REC-domain are conserved. For characterized genes or chemosensory systems, the relevant references are given in Fig. 1B.

Homologous chemotaxis-like systems control phototaxis in other cyanobacterial species as well. Motility in hormogonia of *Nostoc punctiforme* depends on the Hmp system, while the Ptx system is essential for their phototactic orientation (24, 25). In a recently isolated motile *Synechococcus elongatus* strain UTEX 3055, the so-called Pix system was shown to regulate both positive and negative phototaxis (26).

Two additional systems involving CheY-like response regulators control phototactic orientation in *Synechocystis* (Fig. 1A). The two-component regulatory system UirS-UirR triggers negative phototaxis in response to UV light by inducing the expression of the response regulator LsiR (27). The cyanobacteriochrome UirS is also able to sense ethylene, which represses the transcription of *lsiR*, resulting in a more robust positive phototaxis response (28, 29). The photoreceptor PixD interacts with the response regulator PixE to form complexes that dissociate under high-intensity blue-light irradiation (30, 31). Unbound PixE is thought to reverse phototactic orientation and is crucial for the polar assembly of TFP during negative phototaxis under intense blue illumination (3, 32). While the molecular function of LsiR is enigmatic, it was recently established that PixE interacts with the motor ATPase PilB1 (33).

An intriguing feature of these signaling pathways is the specific domain structure of the CheY-like response regulators. The typical cyanobacterial chemotaxis systems each contain two response regulators. While one comprises only the archetypical CheY receiver domain (REC), the other has an N-terminal PATAN domain (which corresponds to DUF4388/PF14332 in the PFAM database) fused to the REC domain (Fig. 1A). In *Synechocystis*, this bipartite domain structure (PATAN-REC) is also found in the response regulators PixE, LsiR, and in a so far uncharacterized CheY-like protein Slr1594 (19). This architecture is the hallmark of PatA-type response regulators (34) named after the *Anabaena* sp. PCC 7120 (*Anabaena*) PatA protein that is required for the accurate patterning of nitrogen-fixing heterocysts along cell filaments (35, 36). PatA localizes to sites of cell division and was recently shown to interact with components of the divisome in *Anabaena* (36, 37).

A detailed characterization of a PATAN-domain has not been published and its molecular function is still unknown. To gain insight into the role of this domain for the function of PatA-type response regulators, we compared the localization and protein interaction of the different PatA-type regulators with their REC-only counterparts (CheY-type from here on) and tested the behavior of mutants expressing isolated PATAN-domains.

## Results

### Phylogeny of PatA-type response regulators

PatA-type response regulators like PixE, PixG, or PatA act as output proteins for unrelated signaling pathways, such as phototaxis and heterocyst formation. We inferred a maximum-likelihood tree build from PATAN-domain containing response regulators in a sample set of cyanobacteria to unravel their phylogeny (Fig. 1). Considering that most detected homologs represent the first gene within a chemotaxis locus, we additionally constructed a phylogenetic tree for these systems based on the CheA sequences (Fig. S1). Wuichet and Zhulin (38) identified four paralogous chemotaxis clusters (Type 1-4 systems) that harbor MCPs with greatly divergent sensory modules within five cyanobacterial genomes. All PatA-type regulators cluster with high confidence into distinct clades that perfectly match the phylogeny of the four evolutionarily conserved chemotaxis systems (Fig. 1B). These systems were classified as chemotaxis systems involved in TFP-based motility (18) and except for Type 4 systems there is multiple evidence that these chemotaxis systems indeed regulate twitching motility in cyanobacteria (see References in Fig 1). Importantly, PatA-type regulators that are not part of a chemosensory system - like the eponymous PatA - do not form a monophyletic cluster but are branching from within different clades. An evaluation of the receiver domains suggests that phosphorylation of the conserved aspartate has been lost in some of these genes (Fig. S2). The phylogenetic analysis thus implies that distinct “orphan” PatA-type regulators evolved independently through gene duplication of chemosensory PatA-type proteins in different cyanobacterial lineages. At least some of them were re-wired to new, phosphorylation-independent sensory pathways as is the case in *Synechocystis* (Fig. 1A). Here the successive duplications of a Tax2 associated gene lead to the emergence of LsiR, PixE, and a third, so far uncharacterized PatA-type regulator encoded by *slr1594*. Given that all other PATAN domain-containing proteins in *Synechocystis* are part of a chemosensory system or are crucial for reversal of phototactic orientation, it is tempting to speculate that Slr1594 might have a similar function. As the gene is co-transcribed with an EAL-domain protein (39) which could function in the degradation of c-di-GMP, Slr1594 might link phototactic orientation to changes in the concentration of this second messenger.

### PatA-type regulators localize to the cytoplasmic membrane

Previous work determined that PixE partially localizes to the cell membrane and interacts with the motor ATPase PilB1 (33). To systematically study the subcellular localization of the different PatA-type regulators in *Synechocystis*, we expressed these as fusion proteins containing a C-terminal enhanced yellow fluorescent protein (eYFP). For comparison, we constructed fluorescent versions of the CheY-like PilH and PixH regulators, encoded in the same *tax* gene cluster but which are missing the PATAN domain (Fig. 1). To avoid any polar effects on the expression of the chemotaxis operons we inserted the expression cassette at a neutral site in the chromosome of the wild-type strain under control of the copper-repressed promoter P*petJ* (40). After verifying that fluorescence is specifically induced in copper-depleted medium (Fig. S3), we analyzed the distribution of the fluorescence signal in cells grown on solid medium using epifluorescence microscopy. Fig. 2 clearly shows that all PatA-type fusion proteins localize to the cell periphery. While the orphan PATAN-CheY regulators show a patchy distribution, the ones encoded from a chemotaxis locus are evenly distributed along the membrane. This is in stark contrast to the PilH-eYFP and PixH-eYFP CheY-type proteins that, accumulate mainly in the cytoplasm, with only a very small fraction being visible at the membrane.

**Fig. 2:**
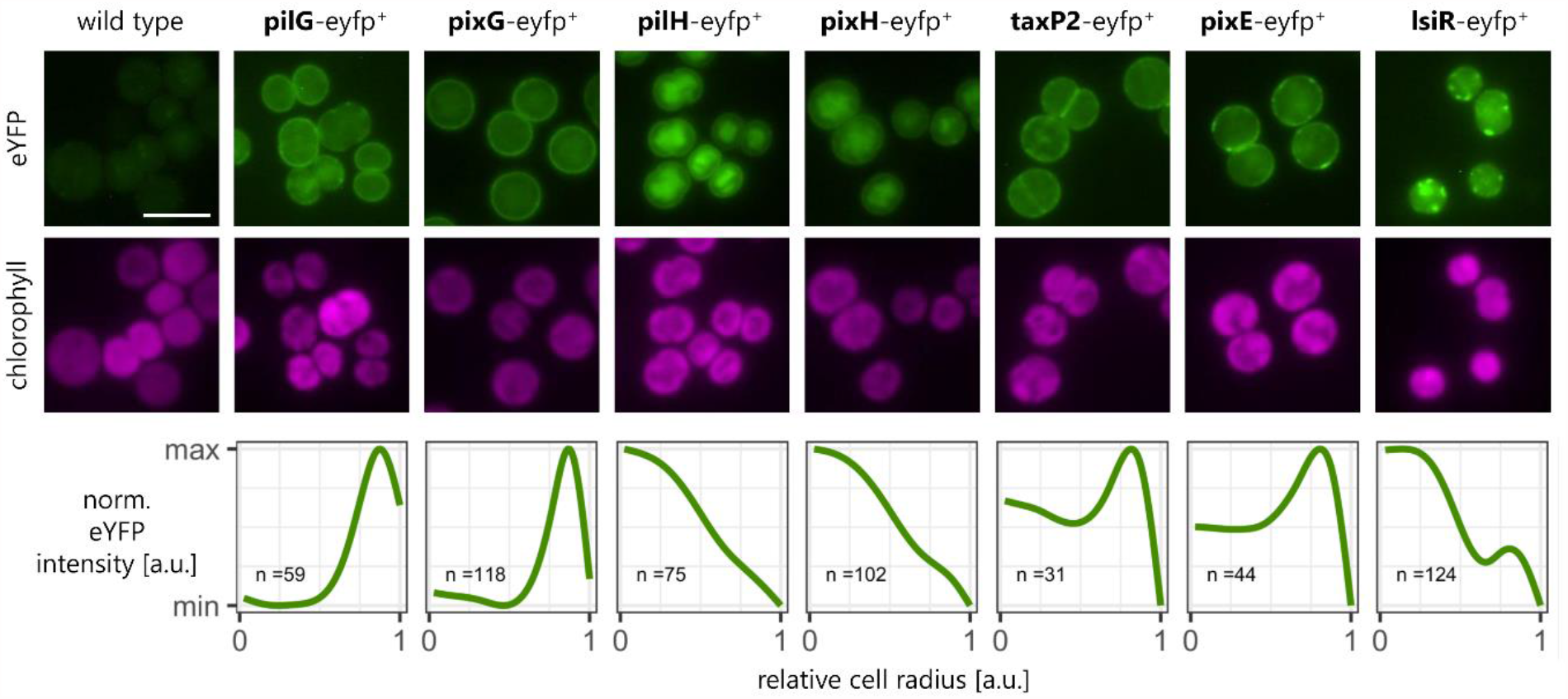
PatA-type regulators localize to the inner membrane. (Upper panel) Fluorescence microscopy images show the subcellular localization of PatA-type and CheY-type response-regulators involved in the regulation of phototaxis, TFP biogenesis, and adhesion. The proteins harboring a C-terminal eYFP fusion were expressed from a neutral genomic locus in a wild-type *Synechocystis* strain. Cells were grown on 0.5% BG11 agar plates without copper to induce gene expression. eYFP fluorescence is shown in green and chlorophyll fluorescence in magenta. Scale bars = 5 µm. (Lower panel) Relative eYFP fluorescence distribution from cell center to cell periphery. The curves were fitted from the radial intensity profiles of n cells and normalized to the WT fluorescence distribution.

### PatA- but not CheY-type regulators interact with the pilus extension motor

Notwithstanding their divergent evolutionary history, regulators like PixG, LsiR, or PixE have a remarkably similar function in regulating phototactic orientation, and we hypothesize that the entire PatA protein family might share a conserved mode of action. Considering the membrane localization and the previously established interaction of PixE with the extension motor PilB1 (33), it seems likely that these proteins directly target components of the motility apparatus. We screened possible interactions of the response regulators PilG/H, PixG/H, PixE, and LsiR that are controlling motility with the essential TFP motor components PilB1, PilT1, and the platform subunit PilC using yeast two-hybrid (Y2H) assays (Fig. 3 and Fig. S4). To include the integral membrane protein PilC in the screen, we generated two constructs to separately express both the N-terminal and C-terminal cytoplasmic domains (PilC_NTD_ residues 1-173 and PilC_CTD_ 239-377, respectively). Additionally, we included PilT2 in the assay, as this motor ATPase has been shown to interact with PilG in a bacterial two-hybrid assay (41).

**Fig. 3:**
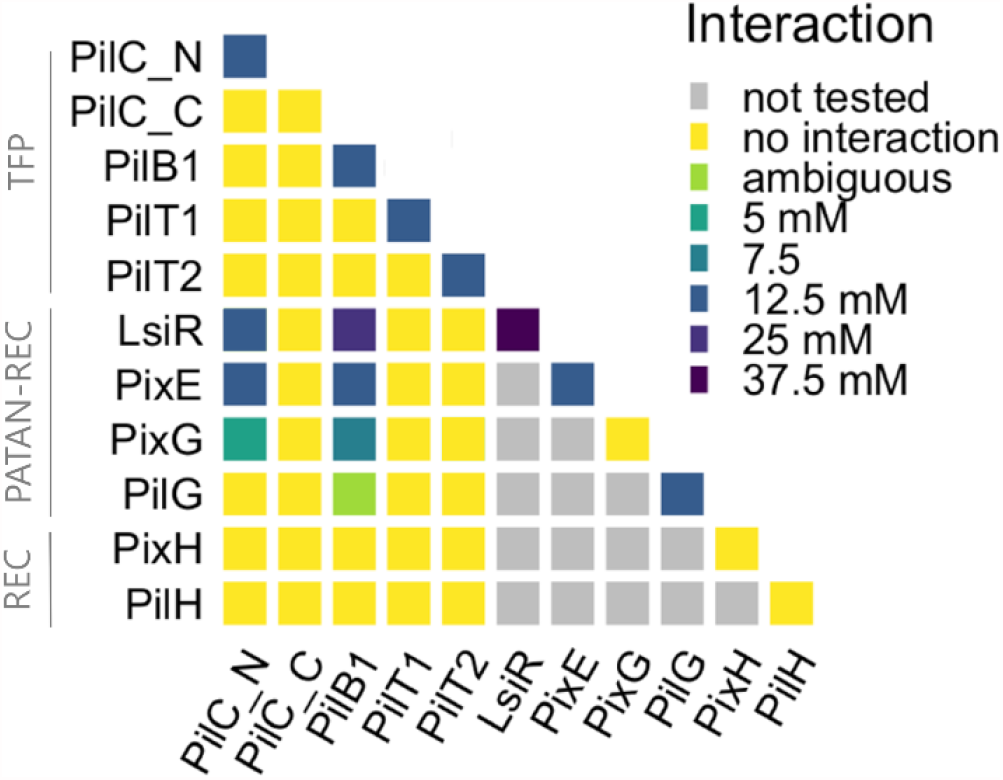
PatA-type regulators interact with the TFP extension motor. Interaction matrix of response regulators and TFP components from the inner membrane. Protein-protein interactions were tested using a Y2H screen by transforming yeast strain AH109 with prey (GAL4 AD) and bait (GAL4 BD) vectors and selection on leucine-, tryptophan-and histidine-depleted selection medium supplemented with different amounts of the competitive inhibitor 3-AT. The highest permissive 3-AT concentration is shown. The PilG-PilB1 is denoted as ambiguous as only four out of six tested yeast clones showed interaction.

Collectively, the Y2H assays established that not only PixE but all tested PatA-type regulators interact with the extension motor ATPase PilB1 but not with the retraction motor ATPase PilT1 nor with PilT2 (We attribute the discrepancy in PilT2 interaction with the relatively small overlap and high false-positive rate observed between different two-hybrid methods). Moreover, we detected interactions between PixE, LsiR, or PixG and the N-terminal cytoplasmic domain of the platform protein (PilC_NTD_). Except for PixG, all PatA-type regulators showed self-interaction, indicative of a possible dimerization. In contrast to the PATAN-domain containing response regulators, the CheY-type proteins PixH and PilH did not interact with any of the pilus proteins and showed no self-interaction in our analysis. Overall, protein interactions of the orphan regulators LsiR and PixE could be detected under more stringent conditions (i.e., higher 3-AT concentrations). It is conceivable that the protein-binding affinity depends on the signaling state of the REC domain, which is most probably phosphorylation-independent in the case of the orphan regulators. On the other hand, the Pix and Pil response regulators, which are part of a chemotaxis system, cannot be phosphorylated by their cognate histidine kinases in the Y2H assays and probably adopt a different signaling state with lower binding affinity.

Apart from the C-terminal domain of PilC, all tested pilus proteins showed self-interaction. Such a multimerization is in agreement with similar results reported previously and indicate that the proteins are correctly folded in yeast. The cytosolic N-terminal domain of *Thermus thermophilus* PilC is known to form a dimer (42) and self-interaction of the putatively hexameric motor ATPases PilB1 and PilT1 has been verified in *Synechocystis* (33, 43). It is noteworthy that no interaction could be observed between the paralogous PilT proteins, indicating that PilT2, which is conserved among many cyanobacteria (10) forms separate homomers. Together, these results suggest that the PATAN-domain containing response regulators might alter TFP extension by binding to PilB1 and the pilus platform PilC.

### The PATAN domain conveys signal output to the TFP

To pinpoint the interacting protein domains, we analyzed the interaction of truncated PixE variants with the PilB1 protein. Similar to full-length PixE, the shortened version containing only the PATAN domain (PixE_PATAN_ residues 1-258) retains its ability to interact with either PilB1 or PilC_NTD_ in the Y2H assay. Conversely, a protein consisting of only the C-terminal PixE receiver domain (PixE_REC_ residues 251-380) did not show any interaction (Fig. 4A). Next, we probed the interaction of full-length PixE, as well as the truncated PixE_PATAN_ variant with different PilB1 constructs to narrow down the interaction surface for the PATAN domain (Fig. 4B). Clearly, the interaction surface lies within the N-terminal domain as only a truncated PilB1_NTD_ residues 1-366) but not the PilB1_CTD_ (residues 367-672) variant showed interaction in YTH assays.

**Fig. 4.**
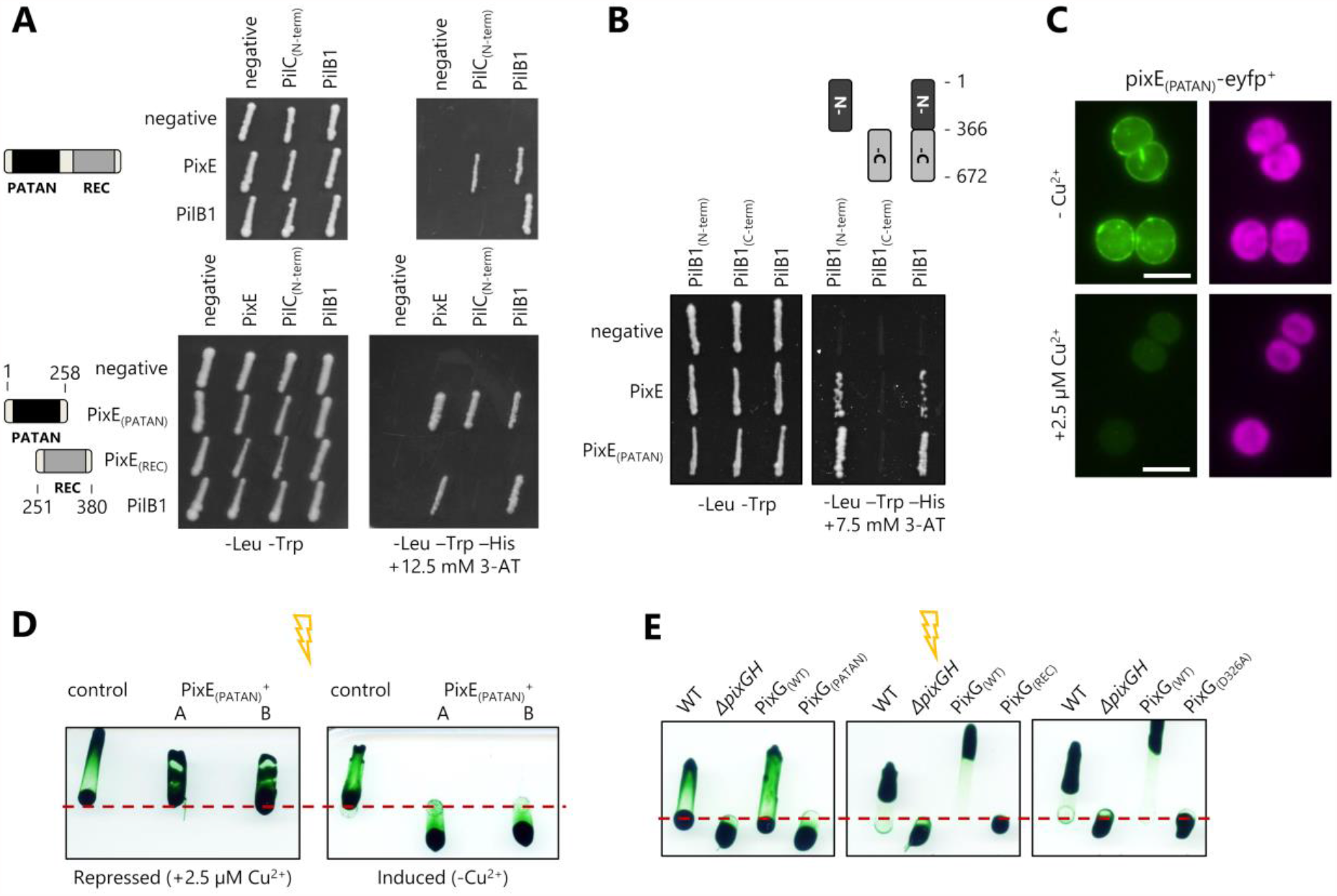
The PATAN domain alone is sufficient to mediate interaction with PilB1 and to switch phototactic orientation. (A) Y2H results show interactions between PixE (upper panel) or truncated versions of PixE (lower panel) with PilB1 or the N-terminal cytoplasmic domain of PilC (PilC_N-term_). For auxotrophic selection, diploid cells harboring prey and bait vectors were grown on restrictive growth medium lacking tryptophan (-Trp), leucine (-Leu). To test specific interaction, plates additionally lacked histidine (-His) and were supplemented with 3-AT. Empty prey and bait vectors (negative) are included as controls. (B) Y2H assays using full-length PixE, the PixE PATAN domain, the N-terminal domain of PilB1, and the C-terminal domain of PilC. (C) Fluorescence microscopy images showing the subcellular localization of the PixE PATAN-domain harboring a C-terminal eYFP fusion. The expression vector and conditions are identical to the strain expressing full-length PixE-eYFP shown in Fig. 2. eYFP fluorescence is shown in green and chlorophyll fluorescence in magenta. Scale bars = 5 µm. (D) Phototaxis assays with *Synechocystis* cells harboring a plasmid for the inducible expression of the PixE PATAN domain (PixE_(PATAN)_^+^) or a control plasmid (control). Cells were spotted onto 0.5% BG-11 motility agar plates and placed in directional white light (∼ 30 μmol photons·m^-2^·s^-1^) indicated by the flash symbol for 11 days. Protein expression was either repressed with 2.5 µM CuSO_4_ or induced by copper depletion. (E) Phototaxis assays with *Synechocystis* wild-type cells, a mutant with a *pixG-pixH* deletion inactivating the Tax1 system (Δ*pixGH*) and complementation strains that encode wild-type *pixH* and either the wild-type *pixG* sequence (PixG_(WT)_) or truncated versions of PixG (PixG_(REC)_ and PixG_(PATAN)_) or a variant with a phosphor-acceptor site mutation (PixG_(D326A)_).

Considering the interactions with PilB1 and PilC_NTD_, we assume that the PATAN-domain localizes the response regulator to the pilus base at the inner membrane. We evaluated the subcellular localization of the PixE PATAN domain which we fused to eYFP using the same strategy as for the full-length PixE. The micrographs in Fig. 4C show a fluorescence distribution along the cell periphery comparable to the full-length protein (see Fig. 2A for comparison), verifying that the N-terminus is sufficient for membrane localization. In order to test whether membrane localization of PatA-type response regulators depends on PilB1 or PilC, we inactivated the genes in *Synechocystis* strains expressing PixG, PilG, TaxP2 and LsiR eYFP-fusion proteins. However, membrane localization of PatA-type response regulators is not lost in *ΔpilB1* or *ΔpilC* mutants (Fig. S5). Thus, we surmise that additional proteins mediate membrane localization of the PatA-type regulators.

Since the Y2H data suggests that the PATAN domain can interact with pilus components independently of other domains, we sought to characterize the role of this interaction during phototaxis in *Synechocystis*. First, we tested phototactic motility in a *Synechocystis* strain transformed with a plasmid encoding the truncated PixE protein under the control of the copper-repressed P*petJ* promoter (PixE_PATAN_^+^). When the ectopic expression of PixE_PATAN_ was repressed, the strain showed positive phototaxis towards white light and was indistinguishable from a *Synechocystis* control strain harboring a control plasmid (Fig. 4D). In stark contrast, PixE_PATAN_^+^ but not the control strain reversed its phototactic orientation and moved away from the light source under conditions that induce protein expression. These experiments imply that PixE_PATAN_itself can bind to the TFP motor also in *Synechocystis* cells and clearly demonstrate that the PixE PATAN domain is sufficient to act as an output signal. In a similar experiment, we analyzed phototaxis in Δ*pixGH* mutant strains complemented with *pixH* and either the wild type *pixG* (PixG_WT_), the truncated variants PixG_PATAN_ or PixG_REC_, or a mutant missing the conserved aspartate at the phosphorylation site (PixG_D326A_). These motility assays (Fig. 4E) demonstrate that only the expression of the full-length PixG with an intact phosphorylation site restored positive phototaxis. This supports the idea that the PatA-type response regulators that are part of chemotaxis operons rely on phosphorylation to switch to an active signaling state.

## Discussion

In *Synechocystis*, action spectra of positive and negative phototaxis overlap with the absorption spectra of photosynthesis (2), and photoreceptor mutants typically reverse orientation but maintain directional movement in regard to the light vector (44). Therefore, it can be reasonably assumed that phototaxis in *Synechocystis* is controlled by two distinct regulatory pathways. The first regulatory circuit establishes cell polarity, presumably through indirect light sensing via alterations in proton motive force (45). At the same time, another system integrates the spectral quality of the incoming light to determine the orientation of polarity and thereby the direction of phototaxis. It is apparently the photoreceptors PixJ1, PixD, and UirS that collectively modulate the decision between positive and negative phototaxis via their cognate PatA-type response regulators. Here we could demonstrate that in *Synechocystis*, the PATAN domain of PatA-type response regulators interacts with the TFP extension motor and that the expression of the PixE PATAN domain alone is sufficient to reverse phototactic orientation. We thus assume that modulation of pilus assembly by the PATAN domain constitutes the principal output of PatA-type regulators in cyanobacteria.

There is direct evidence for an interaction between the cytoplasmic NTD of the platform protein PilC and the motor ATPase PilB (12) and it was suggested that a PilC NTD dimer would fit into the pore formed by a PilB hexamer (14). Given that the PatA-regulators interact with both the cytoplasmic NTD of PilC and the N-terminal domain of PilB1, we speculate that the PATAN domain modulates this interaction between the extension motor and pilus platform. The PATAN domains found in *Synechocystis* and many other bacteria contain a small insertion of two repetitive units, each forming two antiparallel α-helices. Interestingly, repeats of this α-clip domain are also frequently found at the start of the N1 domain of various type II secretion systems (T2SS) and TFP extension ATPases (34). This N1 domain is thought to play a key role in engaging inner membrane components of the TFP machinery (46). Therefore, it is conceivable that the PATAN-domain might share structural similarities with the N-terminus of PilB and that both proteins share - or compete for - a common interaction partner. While we do not know if the self-interaction observed for most PatA-type regulators has any regulatory consequence, we could imagine that this multimer formation might be relevant when many regulators bind the hexameric PilB1 simultaneously.

The REC domain, on the other hand, is likely required to convey sensory input from the MCP through phosphorylation by its cognate histidine kinase. So far, there is no direct evidence of phosphotransfer between CheA homologs and PatA-regulators based on experimental data. In vitro phosphotransfer assays showed that the PatA-type regulator HmpA from *N. punctiforme* does not accept phosphate from the Hmp system (25). However, we observed that the PixG(D326A) variant fails to complement the mutant phenotype (Fig 3E). Together with other circumstantial evidence like the interaction between PilG and the histidine kinase PilL in *Synechocystis* (41), this supports the idea that phosphotransfer is essential for the function of at least some PatA-type regulators from chemosensory systems. The orphan regulators, on the other hand, seem to be mostly phosphorylation independent. The reversal of phototactic orientation in *Synechocystis* expressing only the PATAN domain of PixE (Fig 3D) and the successful complementation of an *Anabaena patA* mutant by a truncated gene missing the REC domain (36) indicate that, in general, the PATAN domain can exert its function independently of other domains.

While we still do not know how the interplay between CheY-type and PatA-type regulators affects the TFP motor, a comparison with chemotactic bacteria suggests a possible mode of action. The disparate localization patterns of the chemosensory response regulators PixG/PixH and PilG/PilH (Fig 2A) are reminiscent of the chemosensory Chp system that regulates twitching migration in *Pseudomonas aeruginosa* (47). *P. aeruginosa* PilG, which does not harbor a PATAN domain, localizes to the cell pole where it promotes polar recruitment of the extension motor PilB favoring forward movement. PilH, which is diffusely localized in the cytoplasm, counteracts unipolar PilB activity, thereby facilitating reversals of cell movement (48). We envision that the response regulators of cyanobacterial Tax systems follow a similar division of labor with the PatA-type regulators promoting unipolar PilB1 activity by directly binding to the TFP motor.

PixE has been shown to co-localize with PilB1 (33) and has been identified in protein complexes up to ∼450 kDa in membrane fractions separated by native PAGE (49). The truncated PixE PATAN domain localizes to the membrane (Fig 3C) indicating that it determines subcellular localization. Nonetheless, membrane localization of different PatA-type regulators is not lost by the deletion of the *pilB1* or *pilC* genes, suggesting that other binding partners exist. Several studies in *Synechocystis* and other cyanobacteria indicate that a variety of proteins that are linked to TFP function can interact with PilB1 among them, HmpF (45, 50), EbsA (51), Hfq (43) and Slr0845 (10). Hence, one must assume that the pilus motor complex is a protein interaction hub where distinct protein subassemblies regulate (specific) TFP with various functions. We speculate that apart from PilB1 and PilC, *Synechocystis* PatA-type regulators interact with other proteins to determine the orientation of pilus assembly during phototaxis. A promising candidate would be HmpF, a filament-forming protein essential for motility in *Synechocystis* and *N. punctiforme* that exhibits dynamic, unipolar localization to the leading cell poles in motile filaments (45, 50, 52, 53).

An identical molecular function for all PatA-type regulators encoded within chemosensory Tax loci is supported by experimental evidence in other cyanobacteria. HmpA, which is part of a Type 1 system (Fig. 1) that regulates motility in *N. punctiforme*, co-localizes with PilA at the junctional pore complexes (25, 52). The regulators PtxA (Type 2) and PixG (Type 4) control directionality during phototactic movement of *N. punctiforme* and *S. elongatus* (24, 26). On the other hand, the orphan regulator PatA localizes to the polar regions of heterocysts or the central plane of dividing cells when overexpressed in *Anabaena* (36, 37). An interaction with the TFP motor during nitrogen starvation or cell division is not immediately plausible. These findings corroborate that at least some orphan PatA-type regulators evolved specific functions other than controlling TFP assembly. Nonetheless, the distinct localization patterns of PatA support the idea that this orphan regulator likewise regulates spatiotemporal protein dynamics. In light of these findings, we argue that a predicted role of cyanobacterial PatA-type regulators as DNA binding proteins (34) seems unlikely.

PatA-type response regulators are typical for cyanobacteria, whereas PATAN-containing proteins of variable domain architectures are widespread in other environmental bacteria. Many genomes encode multiple PATAN-domain-containing proteins (IPR025497/DUF4388), with some species like *M. xanthus* having more than a dozen. While the majority of the more than 8000 proteins containing a DUF4388 annotation comprise no additional domains or harbor a single REC domain, other proteins contain a variety of different domains. Remarkably, there is a number of proteins that are annotated as containing a GspE_N domain (PF05157) which is prototypical for the N-terminus of PilB1. This co-occurrence might hint at an evolutionarily conserved association between the PATAN domain and T2SS assembly ATPases. What little is known from PATAN domain proteins outside the cyanobacteria also hints at a connection between this domain and the TFP apparatus. In *M. xanthus*, the PATAN-domain protein SgnC is required for social gliding motility (54). Furthermore, homologs of MglA and MglB that control asymmetrical assembly of TFP by sorting PilB and PilT to opposite cell poles are regularly encoded in the genetic neighborhood to PATAN-containing proteins (55). It remains to be seen, if there is a conserved role of PATAN-domains in the spatiotemporal regulation of TFP machineries.

## Materials & Methods

### Phylogenetic and sequence analysis

PatA-type response regulators in indicated genomes were detected with hmmsearch using an HMM model generated from an alignment of the six PATAN-containing regulators from *Synechocystis*. Protein sequences were aligned with MAFFT and subsequently refined using Guidance2 (56) with the following settings (bootstraps 100; maxiterate 1000, localpair). Unreliable columns with GUIDANCE scores below 0.93 were discarded. The maximum likelihood phylogenetic tree was inferred using an LG model with a discrete gamma distribution (+G). The percentages at nodes are bootstrap probabilities calculated using 500 replicates. We considered phosphorylation of REC domains unlikely when either the second Aspartate of the conserved pair following β-sheet 1, the Aspartate of the phosphorylation site, or the conserved Threonine/Serine at the end of β-sheet 4 was missing (57) .

### Culture condition and strains

The glucose-tolerant motile *Synechocystis* sp. PCC 6803 PCC-M wild type and mutants (Table S1) were propagated on BG11 agar plates (0.75% (w/v) supplemented with 0.3% (w/v) sodium thiosulfate (Rippka et al., 1979). Liquid cultures were grown in BG11 medium supplemented with 10 mM *N*-[Tris-(hydroxymethyl)-methyl]-2-aminoethanesulphonic acid (TES) buffer (pH 8.0) at 30°C under continuous white-light illumination (Philips TLD Super 80/840) of 50 μmol photons m^-2^ s^-1^ and constant shaking. For induction of the P*petJ* promoter CuSO_4_ was omitted from the medium. For mutant strains antibiotics were added at the following concentrations: chloramphenicol, 7 μg ml^-1^; streptomycin, 10 μg ml^-1^; kanamycin 40 µg ml^-1^; zeocin, 5 μg ml^-1^, gentamycin 10 µg ml^-1^.

### Mutagenesis and plasmid construction

All primers and plasmids are listed in Supplementary Tables S1 and S2. *Synechocystis* strains expressing the PatA-type response regulators with a C-terminal eYFP tag under the control of a copper-sensitive *petJ*-promoter were constructed as follows. The *Synechocystis* genes *pixG* (sll0038), *pixH* (sll0039), *pilG* (slr1041), *lsiR* (slr1214), and *pixE* (slr1693), were amplified from genomic DNA using primers to introduce NdeI and XhoI restriction sites. Each fragment was subsequently cut and ligated into the NdeI and XhoI restriction site of the pSK-hfq-eyfp vector harboring a C-terminal eYFP (Schuergers 2014). The resulting plasmids were verified by sequencing and used to transform wild-type *Synechocystis*, leading to the integration of the expression cassette into a neutral locus via homologous recombination (40). Furthermore, these strains were transformed with genomic DNA from mutant strains to inactivate the *pilB1* or *pilC* genes. For inducible expression in PixE_(PATAN)_^+^ the PixE PATAN domain was amplified from genomic DNA using primers to introduce NdeI and BglII restriction sites and subsequently cut and ligated into the appropriate restriction site of a pUR-expression vector harboring a C-terminal FLAG sequence (pUR-C-FLAG). For the deletion and subsequent complementation of the *pixGH* genes *Synechocystis* cells were transformed with plasmids pUC-ΔpixGH and pUC-pixG(WT)H, which were constructed by seamless cloning. Variants containing a mutated pixG gene were constructed by oligonucleotide-directed mutagenesis of pUC-pixG(WT)H. Complete segregation of these mutants was verified by colony PCR.

### Phototaxis assays

The phototactic movement was analyzed as described previously (58). In short, cell suspensions were spotted on 0.5% (w/v) BG11 agar plates supplemented with 0.2% glucose and 10 mM TES buffer (pH 8.0). For repression of the *petJ* promotor 2.5 µM CuSO_4_ was added. After 2–3 days under diffuse white light (50 μmol photons m^-2^ s^-1^) the plates were placed into non-transparent boxes with a one-sided opening (> 5 μmol photons m^-2^ s^-1^) for at least 10 days.

### YTH analysis

The YTH analysis was carried out as described previously (33). Primers for the construction of yeast expression plasmids are listed in Supplementary Table S1. *Synechocystis* DNA fragments were amplified from genomic DNA, subsequently cut with the appropriate restriction enzymes, and ligated into the appropriately cut pCGADT7ah, pGADT7ah, pD153, or pGBKT7 bait- and prey-vectors, creating either N- or C-terminally tagged GAL4 activation domain (AD/prey) or GAL4 DNA-binding domain (BD/bait) fusion proteins. Yeast transformants containing bait-prey pairs were selected on complete supplement mixture (CSM) dropout medium lacking leucine and tryptophan at 30°C for 4 days. Subsequently, cells were screened for interaction by streaking on dropout medium lacking leucine, tryptophan, and histidine supplemented with different amounts of 3-amino-1,2,4-triazole (3-AT) at 30°C for 6-7 days.

### Epifluorescence microscopy

Micrographs were captured at room temperature using an upright Nikon Eclipse Ni-U microscope fitted with an x40 (numerical aperture 0.75) and an x100 oil immersion objective (NA 1.45). For visualization of epifluorescence, a GFP-filter block (excitation 450–490 nm; emission 500–550 nm; exposure time 1 s) or a Cy3-filter block (excitation 530–560 nm; emission 575–645 nm; exposure time <50 ms) were used. Cells were picked from colonies grown on phototaxis plates, resuspended in a small volume of PBS pH 7.4 buffer, and spotted on glass slides. Fluorescence distribution was analyzed with ImageJ (1.53c) using the Radial Profile Extended plugin and normalization and curve fitting (LOESS) was calculated using custom R code.

## Acknowledgments

We would like to thank Conrad W. Mullineaux (Queen Mary University of London) for helpful discussions and preliminary work that could not be included in the final manuscript and Thomas Wallner (University of Freiburg) for providing the pUR-C-FLAG plasmid. This work was supported by grants to AW in frame of the SFB 1381 by the Deutsche Forschungsgemeinschaft (German Research Foundation) under project no. 403222702-SFB 1381 (A2) and under project no. Wi 2014/8-1. YH was supported by the China Scholarship Council (CSC).

