## Supplementary Information for "PATAN-domain response regulators interact with the Type IV pilus motor to control phototactic orientation in the cyanobacterium *Synechocystis* sp. PCC 6803"

*corresponding author

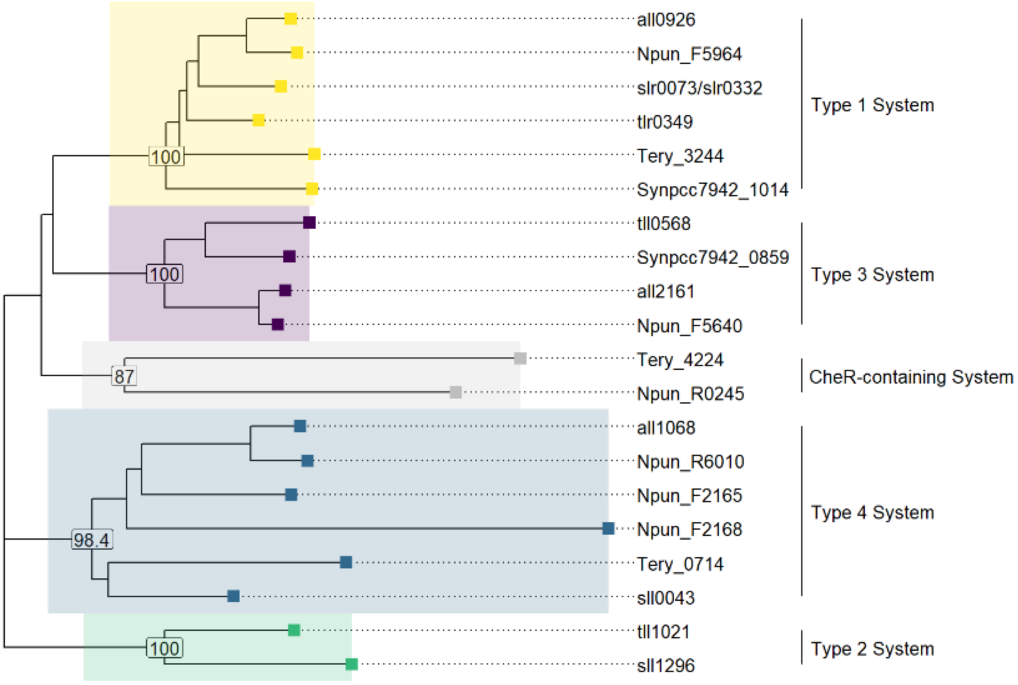


**Figure S1: CheA Histidine-Kinases in cyanobacteria are part of distinct chemosensory systems**. The maximum likelihood phylogenetic tree was constructed from CheA homologs found in the chemosensory systems of a subset of cyanobacteria listed in the MiST3.0 database (Gumerov et al., 2020). Sequences were aligned with MAFFT and unreliable columns removed by GUIDANCE2 (--bootstraps 100--maxiterate 1000 –localpair) (Sela et al., 2015). The evolutionary history was inferred using an LG model (Le and Gascuel, 2008) with a discrete gamma distribution (+G). The percentages at nodes are bootstrap probabilities calculated using 500 replicates. For clarity, the N- and C-terminal PilL sequences from *Synechocystis* (encoded by two annotated open reading frames *slr0073* and *slr0322*) were concatenated before the analysis.

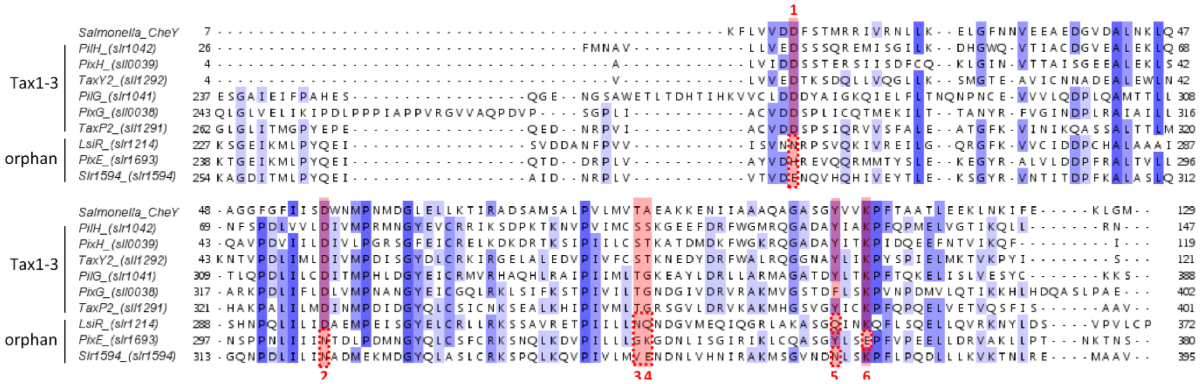

**Figure S2: REC domain conservation of CheY- and PatA-type response regulators.** Multiple sequence alignment of the REC domains from the *Synechocystis* sp. PCC6803 CheY- and PatA-type proteins compared to *Salmonella* CheY. Sequence identity is shown in blue and columns implicated in protein phosphorylation are shaded red. (1) D critical for metal ion binding; (2) conserved phosphor-accepting D; (3) T/S interact with phosphoryl group; (4) typically A/G but sometimes S/T: allows access to phosphorylation site; (5-6) (F/Y)xxK motif important for phosphorylation-mediated conformational changes.

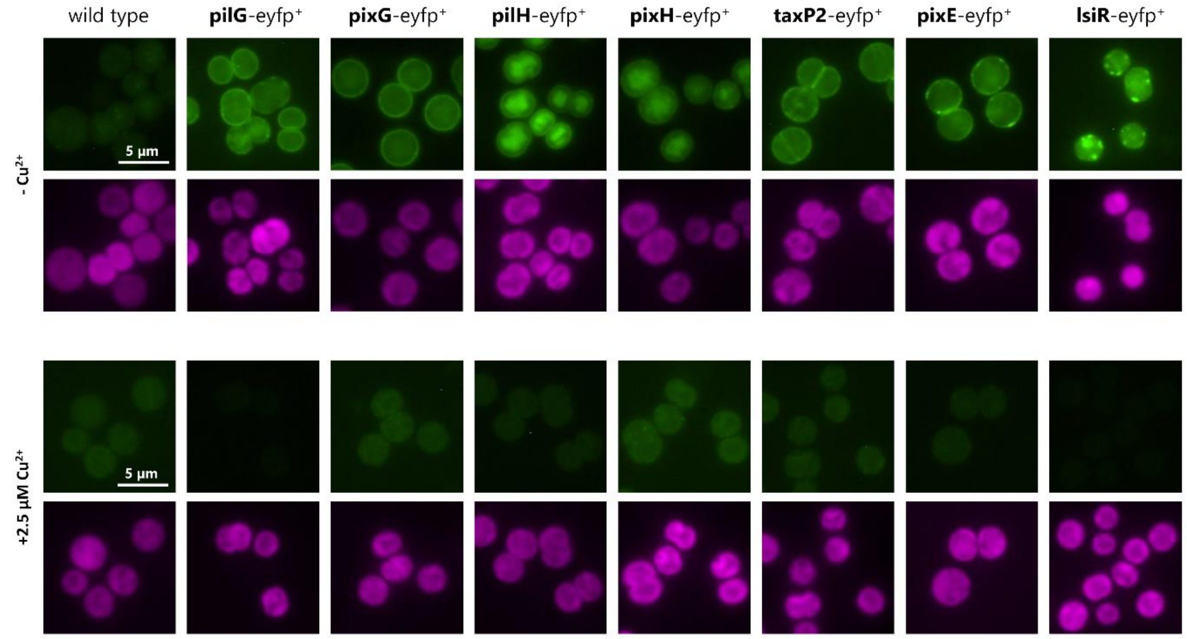


**Figure S3: Inducible expression of fluorescence-tagged response regulator proteins***.* The proteins harboring a C-terminal eYFP fusion are expressed under the control of the copper-repressed *petJ* promoter. The expression cassette was inserted into a neutral genomic locus in wild-type *Synechocystis* cells via homologous recombination. Cells were grown on 0.5% BG11 agar plates without copper to induce gene expression or with the addition of 2.5 µM $CuSO_4$ to repress the *petJ* promoter. eYFP fluorescence is shown in green and chlorophyll fluorescence in magenta. Scale bars = 5 µm.

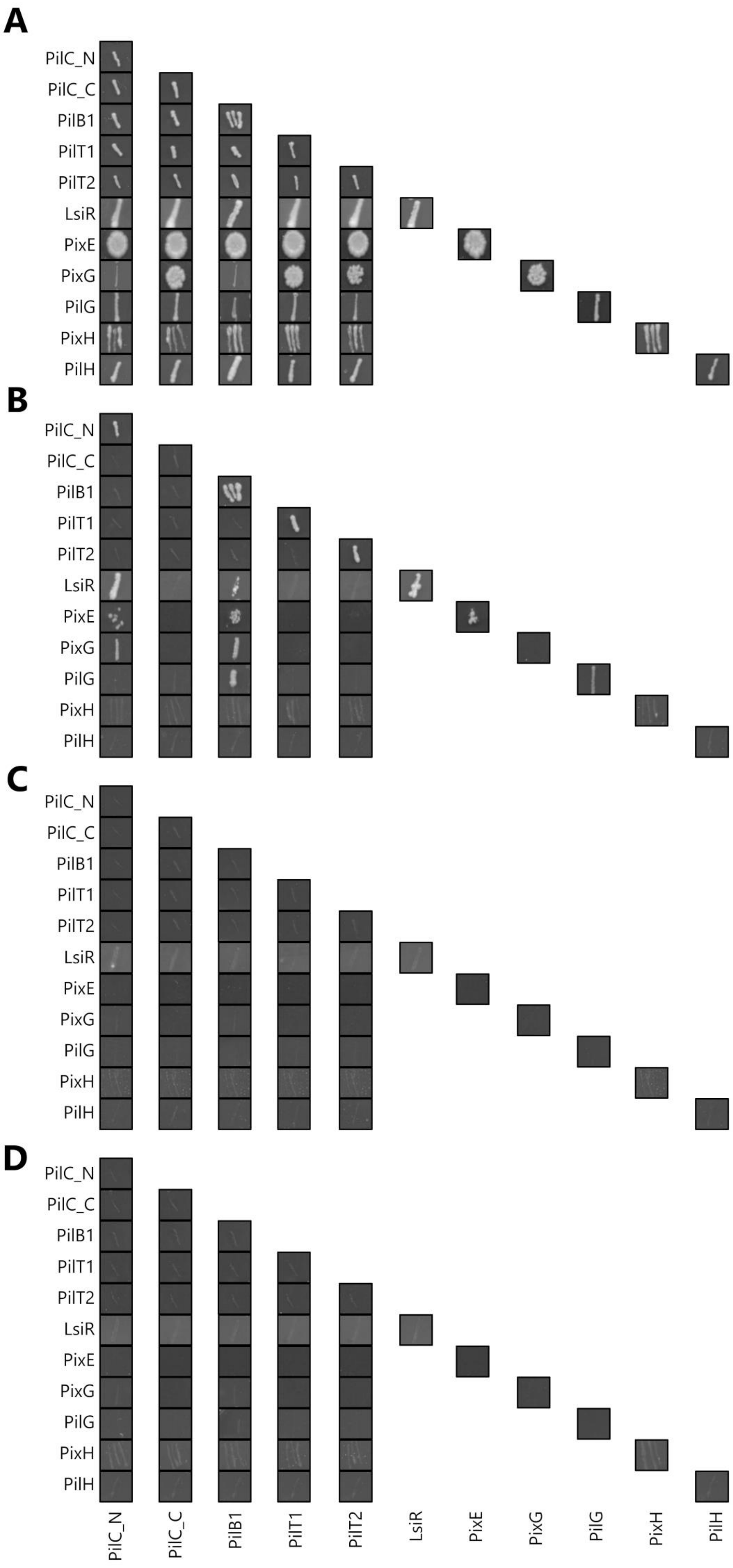
A
B
C
D
PilC_N
PilC_C
PilB1
PilT1
PilT2
LsiR
PixE
PixG
PilG
PixH
PilH

**Figure S4: Y2H analysis.** Protein-protein interactions were tested using a Y2H screen by transforming yeast strain AH109 with prey (GAL4 activation domain) and bait (GAL4 DNA binding domain) vectors and selection on complete supplement mixture dropout medium supplemented with different amounts of the competitive inhibitor 3-AT. Representative results using the highest permissive 3-AT concentration are shown (for 3-AT concentrations see Fig. 3 in the main text). (A) Yeast-growth after 6-7 days at 30°C on (A) permissive medium (-Leu/-Trp) and (B) selective medium (-Leu/-Trp/-His). (C-D) *bait*- and *prey*-only controls on selective medium.

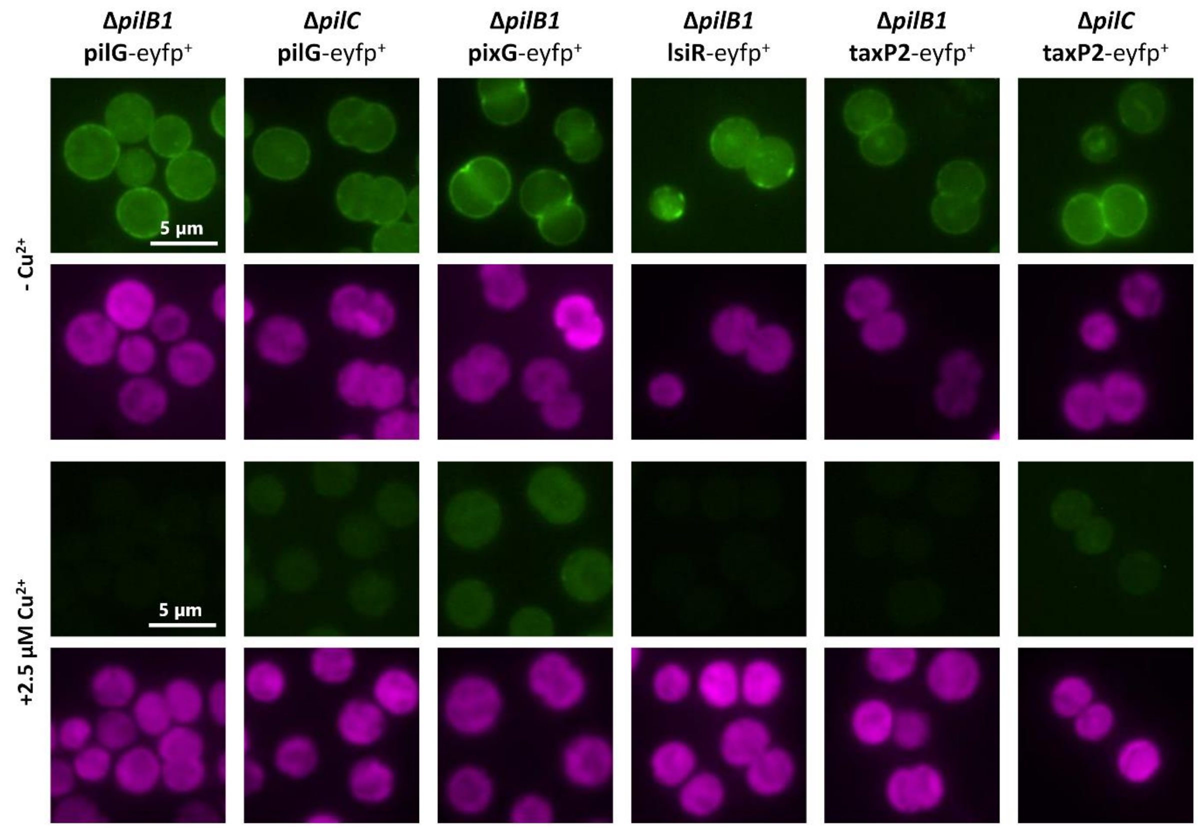


**Figure S5: Localization of PatA-type response regulators in Δ*pilB1 or ΔpilC* mutant strains.** The proteins harboring a C-terminal eYFP fusion are expressed under the control of the copper-repressed *petJ* promoter. The expression cassette was inserted into a neutral genomic locus in wild-type *Synechocystis* cells via homologous recombination. Cells were grown on 0.5% BG11 agar plates without copper to induce gene expression (left) or with the addition of 2.5 µM $CuSO_4$ to repress the *petJ* promoter (right). eYFP fluorescence is shown in green and chlorophyll fluorescence in magenta. Scale bars = 5 µm.

### Table S1. Strains and Plasmids used in this study.

| Bacterium/plasmids | Relevant characteristic(s) | Reference/Source |
|---|---|---|
| ***Synechocystis* strains** | | |
| *Synechocystis* WT | PCC-M wild type | (Trautmann et al., 2012) |
| pixE$_{(PATAN)}$$^{+}$ | WT containing plasmid pUR-*pixE_PATAN_*flag | This study |
| pixE$_{(PATAN)}$-eyfp$^{+}$ | WT transformed with pSK-*pixE_PATAN_*eyfp | This study |
| pilG-eyfp$^{+}$ | WT transformed with pSK-*pilG*-eyfp | This study |
| pixG-eyfp$^{+}$ | WT transformed with pSK-*pixG-eyfp* | This study |
| pilH-eyfp$^{+}$ | WT transformed with pSK-*pilH-eyfp* | This study |
| pixH-eyfp$^{+}$ | WT transformed with pSK-*pixH-eyfp* | This study |
| taxP2-eyfp$^{+}$ | WT transformed with pSK-*taxP2-eyfp* | This study |
| pixE-eyfp$^{+}$ | WT transformed with pSK-*pixE-eyfp* | This study |
| lsiR-eyfp$^{+}$ | WT transformed with pSK-*lsiR-eyfp* | This study |
| Δ*pilB1* | *pilB1* knockout mutant (Zeo$^{R}$) | (Linhartová et al., 2014) |
| Δ*pilC* | *pilC* knockout mutant (Str$^{R}$) | (Bhaya et al., 2000) |
| Δ*pilB1*/lsiR-eyfp$^{+}$ | lsiR-eyfp$^{+}$ transformed with genomic DNA of *ΔpilB1* | This study |
| Δ*pilB1*/pixG-eyfp$^{+}$ | pixG-eyfp$^{+}$ transformed with genomic DNA of *ΔpilB1* | This study |
| Δ*pilB1*/pilG-eyfp$^{+}$ | pilG-eyfp$^{+}$ transformed with genomic DNA of *ΔpilB1* | This study |
| Δ*pilB1*/taxP2-eyfp$^{+}$ | taxP2-eyfp$^{+}$ transformed with genomic DNA of *ΔpilB1* | This study |
| Δ*pilC*/pilG-eyfp$^{+}$ | pilG-eyfp$^{+}$ transformed with genomic DNA of *ΔpilC* | This study |
| Δ*pilC*/taxP2-eyfp$^{+}$ | taxP2-eyfp$^{+}$ transformed with genomic DNA of *ΔpilC* | This study |
| *ΔpixGH* | *Synechocystis* transformed with pUC-ΔpixGH | This study |
| PixG$_{(WT)}$ | *ΔpixGH* transformed with pUC-pixG(WT)H | This study |
| PixG$_{(PATAN)}$ | *ΔpixGH* transformed with pUC-pixG(PATAN)H | This study |
| PixG$_{(REC)}$ | *ΔpixGH* transformed with pUC-pixG(REC)H | This study |
| PixG$_{(D326A)}$ | *ΔpixGH* transformed with pUC-pixG(D326A)H | This study |
| **Y2H plasmids** | | |
| pGADT7ah | Expression vector in yeast cells; N_GAL4 AD, *LEU2*; Amp$^{R}$ | (Hiltbrunner et al., 2005) |
| pCGADT7ah | Expression vector in yeast cells; C_GAL4 AD, *LEU2*; Amp$^{R}$ | (Rausenberger et al., 2011) |
| pGBKT7 | Expression vector in yeast cells; N_GAL4 BD, *TRP1*; Km$^{R}$ | Clontech |
| pD153 | Expression vector in yeast cells; C_GAL4 BD, *TRP1*; Amp$^{R}$ | (Shimizu-Sato et al., 2002) |
| pGAD-*pilC_N* | AD-*pilC_N* cloned into pGADT7ah | This study |
| pGAD-*pilC_C* | AD-*pilC_C* cloned into pGADT7ah | This study |
| pGAD-*pilT2* | AD-*pilT2* cloned into pGADT7ah | This study |
| pGAD-*lsiR* | AD-*lsiR* cloned into pGADT7ah | This study |
| pGAD-*pilG* | AD-*pilG* cloned into pGADT7ah | This study |
| pGAD-*pilH* | AD-*pilH* cloned into pGADT7ah | This study |
| pGAD-*pilB1_N* | AD-*pilB1_N* cloned into pGADT7ah | This study |
| pGAD-*pilB1_C* | AD-*pilB1_C* cloned into pGADT7ah | This study |
| pCGAD-*pilC_N* | *pilC_N*-AD cloned into pCGADT7ah | This study |
| pCGAD-*pilC_C* | *pilC_C*-AD cloned into pCGADT7ah | This study |
| pCGAD-*pilT2* | *pilT2*-AD cloned into pCGADT7ah | This study |
| pCGAD-*lsiR* | *lsiR*-AD cloned into pCGADT7ah | This study |
| pCGAD-*pilG* | *pilG*-AD cloned into pCGADT7ah | This study |
| pCGAD-*pilH* | *pilH*-AD cloned into pCGADT7ah | This study |
| pCGAD-*pilB1_N* | *pilB1_N*-AD cloned into pCGADT7ah | This study |
| pCGAD-*pilB1_C* | *pilB1_C*-AD cloned into pCGADT7ah | This study |
| pGBK-*pilC_N* | BD-*pilC_N*cloned into pGBKT7 | This study |
| pGBK-*pilC_C* | BD-*pilC_C* cloned into pGBKT7 | This study |
| pGBK-*pilT2* | BD-*pilT2* cloned into pGBKT7 | This study |
| pGBK-*lsiR* | BD-*lsiR* cloned into pGBKT7 | This study |
| pGBK-*pilG* | BD-*pilG* cloned into pGBKT7 | This study |
| pGBK-*pilH* | BD-*pilH* cloned into pGBKT7 | This study |
| pGBK-*pixE-PATAN* | BD-*pixE-PATAN* cloned into pGBKT7 | This study |
| pGBK-*pixE-REC* | BD-*pixE-REC* cloned into pGBKT7 | This study |
| pD153-*pilC_N* | *pilC_N-BD* cloned into pD153 | This study |
| pD153-*pilC_C* | *pilC_C-BD* cloned into pD153 | This study |

| | | |
|---|---|---|
| pD153-*pilT2* | *pilT2-BD* cloned into pD153 | This study |
| pD153-*lsiR* | *lsiR-BD* cloned into pD153 | This study |
| pD153-*pilG* | *pilG-BD* cloned into pD153 | This study |
| pD153-*pilH* | *pilH-BD* cloned into pD153 | This study |
| **Plasmids for the transformation of cyanobacteria** | | |
| pSK-hfq-eyfp | pSDC01-derived vector; $P_{petJ}$, C_eYFP, $Ter_{OOP}$, $Cm^R$ | (Schuergers et al., 2014) |
| pSK-*pixE_PATAN-eyfp* | *pixE-PATAN*-eyfp cloned into pSK-hfq-eyfp, $Cm^R$ | This study |
| pSK-*pilG-eyfp* | *pilG-eyfp* cloned into pSK-hfq-eyfp, $Cm^R$ | This study |
| pSK-*pixG-eyfp* | *pixG-eyfp* cloned into pSK-hfq-eyfp, $Cm^R$ | This study |
| pSK-*pilH-eyfp* | *pilH-eyfp* cloned into pSK-hfq-eyfp, $Cm^R$ | This study |
| pSK-*pixH-eyfp* | *pixH-eyfp* cloned into pSK-hfq-eyfp, $Cm^R$ | This study |
| pSK-*taxP2-eyfp* | *taxP2-eyfp* cloned into pSK-hfq-eyfp, $Cm^R$ | This study |
| pSK-*pixE-eyfp* | *pixE-eyfp* cloned into pSK-hfq-eyfp, $Cm^R$ | This study |
| pSK-*lsiR-eyfp* | *lsiR-eyfp* cloned into pSK-hfq-eyfp, $Cm^R$ | This study |
| pUR-C_flag | Expression vector; $P_{petJ}$, C_FLAG, $Ter_{OOP}$; $Km^R$, $Strep^R$ | T. Wallner (Uni Freiburg) |
| pUR-*pixE_PATAN*_flag | *pixE-PATAN*_flag cloned into pUR-C_flag, $Km^R$, $Strep^R$ | This study |
| pUC-ΔpixGH | pUC19-based *pixGH* KO construct containing a kanamycin-resistance cassette flanked by the 650 bp regions upstream of position -285 and downstream of of position +1706 from the translation start site of *pixG*; $Km^R$, $Amp^R$ | This study |
| pUC-pixG(WT)H | pUC-ΔpixGH based vector for the complementation of *pixGH* where the kanamycin resistence cassette is replaced by a gentamycin-resistance cassette flanked by the +36 to -150 region from TSS of *slr0031* and the -285 to +1706 region from the TSS of *pixG*; $Gen^R$, $Amp^R$ | This study |
| pUC-pixG(PATAN)H | pUC-pixG(WT)H derivative replacing full-length *pixG* with *pixG-PATAN*; $Gen^R$, $Amp^R$ | This study |
| pUC-pixG(REC)H | pUC-pixG(WT)H derivative replacing full-length *pixG* with *pixG-REC*; $Gen^R$, $Amp^R$ | This study |
| pUC-pixG(D326A)H | pUC-pixG(WT)H derivative replacing *pixG* with *pixG-D326A*; $Gen^R$, $Amp^R$ | This study |

### Table S2. Primers used in this study

| Name | Sequence (5′ → 3′) | PCR product |
|---|---|---|
| **RE cloning of Y2H plasmids** | | |
| AD-pilC_N-fw | TACATATGGCTACGTTTGTCGCTC | AD-*pilC_N* |
| AD-pilC_N-rev | TTCTCGAGTTACACCGGATAAGCCATGG | |
| AD-pilC_C-fw | TTCATATGAAAAAATATTACGGAACCTATGC | AD-*pilC_C* |
| AD-pilC_C-rev | CTCGAGTTACATAGCTGGTTCTATAATAC | |
| AD-pilT2-fw | TAGGATCCTGAACCAACCTCCCCGC | AD-*pilT2* |
| AD-pilT2-rev | TTCTCGAGTTAGGTTCTGCCCCGCAG | |
| AD-LsiR-fw | TAGGATCCTGACTGCTGTGATCACCCG | AD-*lsiR* |
| AD-LsiR-rev | TTCTCGAGCTAGGGACAAAGAACAG | |
| AD-pilG-fw | AGATCTATCAGGGAACCCTGAAC | AD-*pilG* |
| AD-pilG-rev | CTCGAGTGATTTTTTACAGTAAGATTCAAC | |
| AD-pilH-fw | AGATCTATATGGAAAATAAACCAGG | AD-*pilH* |
| AD-pilH-rev | CTCGAGATTGCGCAGGAGTTGTTTG | |
| AD-PilB1-fw | GCAGATCTTGACATCTTCCTCCTC | AD-*pilB1_N* |
| AD-pilB1(1-366)-rev | ATCTCGAGCCGGGCGGCCAATTCCCTTAC | |
| AD-pilB1(367-672)-fw | GCAGATCTTGCCCTATGGCTTAATGTTGG | AD-*pilB1_C* |
| AD-PilB1-rev | ATCTCGAGGCTAAACCGGGAAG | |
| pilC_N-BD-fw | TAGGATCCATGGCTACGTTTGTCGCTC | *pilC_N*-AD |
| pilC_N-AD-rev | CGTCTAGACACCGGATAAGCCATGGCG | |
| pilC_C-BD-fw | TAGGATCCATGAAAAAATATTACGGAACCTATGC | *pilC_C*-AD |
| pilC_C-AD-rev | CGTCTAGACATAGCTGGTTCTATAATAC | |
| pilT2-AD-fw | TAGGATCCATGAACCAACCTCCCCGC | *pilT2*-AD |
| pilT2-AD-rev | CGTCTAGAGGTTCTGCCCCGCAGTCG | |
| LsiR-AD-fw | TAAGATCTATGACTGCTGTGATCAC | *lsiR*-AD |
| LsiR-AD-rev | TATCTAGAGGGACAAAGAACAGGGAC | |
| pilG-AD-fw | AGATCTATGCAGGGAACCCTGAAC | *pilG*-AD |
| pilG-AD-rev | TCTAGATGATTTTTTACAGTAAGATTCAAC | |
| pilH-AD-fw | AGATCTATGATGGAAAATAAACCAGGC | *pilH*-AD |
| pilH-AD-rev | TCTAGAATTGCGCAGGAGTTGTTTG | |
| pilB1-AD-fw | GCAGATCTATGACATCTTCCTCCTC | *pilB1_N*-AD |
| pilB1(1-366)-AD-rev | ATGCTAGCCCGGGCGGCCAATTCCCTTAC | |
| pilB1(367-672)-AD-fw | GCAGATCTATGCCCTATGGCTTAATGTTGG | *pilB1_C*-AD |
| PilB1-AD-rev | ATGCTAGCGCTAAACCGGGAAG | |
| BD-pilC_N-fw | TAGGATCCGGCTACGTTTGTCGCTCAAG | BD-*pilC_N* |
| BD-pilC_N-rev | GCACTAGTCACCGGATAAGCCATGGCG | |
| BD-pilC_C-fw | TAGGATCCGAAAAAATATTACGGAACCTATGC | BD-*pilC_C* |
| BD-pilC_C-rev | GAGTCGACTTACATAGCTGGTTCTATAATAC | |
| BD-pilT2-fw | TAGGATCCGAACCAACCTCCCCGC | BD-*pilT2* |
| BD-pilT2-rev | GCACTAGTGGTTCTGCCCCGCAGTCG | |
| BD-LsiR-fw | TAGGATCCGACTGCTGTGATCACCCG | BD-*LsiR* |
| BD-LsiR-rev | GCACTAGTGGGACAAAGAACAGGGAC | |
| BD-pilG-fw | AGATCTACAGGGAACCCTGAACGAAATTG | BD-*pilG* |
| BD-pilG-rev | ACTAGTTGATTTTTTACAGTAAGATTCAAC | |
| BD-pilH-fw | AGATCTAATGGAAAATAAACCAG | BD-*pilH* |
| BD-pilH-rev | ACTAGTATTGCGCAGGAGTTGTTTG | |
| BD-pixE-fw | TAGGATCCAAGCAATTCAGTTTTGTCCAC | BD-*pixE-PATAN* |
| BD-pixE-PATAN-rev | GCACTAGTCACCAAAGGGCGGTCATC | |
| BD-pixE-REC-fw | TAGGATCCACAAACGGATGACCGCCCTTTG | BD-*pixE-REC* |
| BD-pixE-rev | GCACTAGTGGAGTTGGTTTTATTGGTGG | |
| pilC_N-BD-fw | TAGGATCCATGGCTACGTTTGTCGCTC | *pilC_N*-BD |
| pilC_N-BD-rev | GAGTCGACAACACCGGATAAGCCATGGCG | |
| pilC_C-BD-fw | TAGGATCCATGAAAAAATATTACGGAACCTATGC | *pilC_C*-BD |
| pilC_C-BD-rev | GAGTCGACAACATAGCTGGTTCTATAATAC | |
| pilT2-AD-fw | TAGGATCCATGAACCAACCTCCCCGC | *pilT2*-BD |
| pilT2-BD-rev | AAGTCGACAAGGTTCTGCCCCGCAGTCG | |

| | | |
|---|---|---|
| LsiR-AD-fw | TAAGATCTATGACTGCTGTGATCAC | *lsiR*-BD |
| LsiR-BD-rev | CCCGGGACGGGACAAAGAACAGGGAC | |
| pilG-AD-fw | AGATCTATGCAGGGAACCCTGAAC | *pilG*-BD |
| pilG-BD-rev | CCCGGGTTTGATTTTTTACAGTAAGATTCAAC | |
| pilH-AD-fw | AGATCTATGATGGAAAATAAACCAGGC | *pilH*-BD |
| pilH-BD-rev | CCCGGGTTATTGCGCAGGAGTTGTTTG | |
| **RE cloning of plasmids for expression of fusion proteins** | | |
| NdeI-pilG-fw | TGCATATGCAGGGAACCCTG | *pilG*_eyfp |
| Xhoi-pilG-rev | GACTCGAGTGATTTTTTACAGTAAGATTC | |
| NdeI-pixG-fw | TGCATATGACAGCTCCCACCCCCT | *pixG*_eyfp |
| XhoI-pixG-rev | GACTCGAGTTCCGCTGGCAGCGATGC | |
| NdeI-pilH-fw | TGCATATGATGGAAAATAAACCAGGC | *pilH*_eyfp |
| XhoI-pilH-rev | GACTCGAGATTGCGCAGGAGTTGTTTG | |
| NdeI-pixH-fw | TGCATATGGGCAGCGCACTTGTTA | *pixH*_eyfp |
| XhoI-pixH-rev | GACTCGAGGATAAATTGCTTGATTACCGTG | |
| NdeI-taxP2-fw | TGCATATGCAATCTCCCCTGTC | *taxP2*_eyfp |
| XhoI-taxP2-rev | GACTCGAGAACAGCTGCGGAGATAAAG | |
| NdeI-pixE-fw | TGCATATGAGCAATTCAGTTTTGTC | *pixE*_eyfp |
| XhoI-pixE-rev | GACTCGAGGGAGTTGGTTTTATTGGTG | |
| NdeI-lsiR-fw | TGCATATGACTGCTGTGATCACCCG | *lsiR*_eyfp |
| XhoI-lsiR-rev | GACTCGAGGGGACAAAGAACAGGGAC | |
| pixE-PATAN-fw | AAAACATATGAGCAATTCAGTTTTGTCCAC | *pixE_PATAN*_flag |
| pixE-PATAN-rev | AAAAAGATCTCACCAAAGGGCGGTCATC | |
| NdeI-pixE-fw | TGCATATGAGCAATTCAGTTTTGTC | *pixE_PATAN*_eyfp |
| pixE-PATAN-eyfp-rev | AAAACTCGAGCACCAAAGGGCGGTCATC | |
| **Seamless cloning of pUC-ΔpixGH** | | |
| pUC19+upward-fw | TTCGAGCTCGGTACCCATACATTACCCCTGGAGG | upstream homology region |
| Upward+kana-rev | CACGAGGCAGACCTCAACTTTATATCCCCCATGC | |
| Upward+kana-fw | ATGGGGGATATAAAGTTGAGGTCTGCCTCGTGAAG | Kanamycin cassette |
| Kana+downward-rev | GGGGCGTTGGCATCGCTGCAAAAGCCACGTTGTG | |
| Kana+downward-fw | CAACGTGGCTTTTGCAGCGATGCCAACGCCCCAG | downstream homology |
| Downward+pUC19-rev | ACTCTAGAGGATCCCCTAGGGTAAATTCCCAAATAGTCC | region |
| **Seamless cloning of pUC-pixG(WT)H** | | |
| pUC19+upward-fw | TTCGAGCTCGGTACCCATACATTACCCCTGGAGG | upstream homology region |
| Upward+pixH-rev | CAAGCAATTTATCTGAACTTTATATCCCCCATGC | |
| Upward+pixH-fw | ATGGGGGATATAAAGTTCAGATAAATTGCTTGATTACCG | *pixGH* (-285 to +1706 region |
| pixG+gen-rev | TTCGAGCTCGGTACCCCACTTGGTAGTGCAGATG | from *pixG TSS*) |
| pixG+gen-fw | TCTGCACTACCAAGTGGGGTACCGAGCTCGAATTG | Gentamycin cassette |
| Gen+slr0031-rev | CCCCTCGGTGAGGTCGGGGTACCGAGCTCGAATTG | |
| Gen+slr0031-fw | TTCGAGCTCGGTACCCCGACCTCACCGAGGGGAC | *Slr0031* (+36 to -150 region |
| Slr0031+downward-rev | GGGGCGTTGGCATCGCCACTTGGTAGTGCAGATG | from TSS) |
| Slr0031+downward-fw | TCTGCACTACCAAGTGGCGATGCCAACGCCCCAG | downstream homology |
| Downward+pUC19-rev | ACTCTAGAGGATCCCCTAGGGTAAATTCCCAAATAGTCC | region |
| ***pixG* mutagenesis in pUC-pixG(WT)H** | | |
| pixG_PATAN-fw | GATCAGGGGACCACTGGGCAC | *pixG_PATAN* |
| pixG_PATAN-rev | TAACTTGACCCAAGGAGGCC | |
| pixG_REC-fw | CATTTAATCAAAGGAAGAGGCAG | *pixG_REC* |
| pixG_REC-rev | GATGTGCCCAGTGGTCCCCTG | |
| pixG_D326A-fw | GATTTTCCTCGCTTTGGTCATGCC | *pixG_D326A* |
